# Transcriptome Analysis of juvenile Pacific White Shrimp (*Litopenaeus vannamei*) with symptom of stunted growth

**DOI:** 10.1101/546770

**Authors:** Huaiyi Fang, Jingjing Song, Bin Gong, Tingcai Pang, Chunyan Peng

## Abstract

In search for viruses associated with *Litopenaeus vannamei* with symptom of stunted growth, we have undertaken a comparative transcriptome analysis of total RNA sequences using Illumina based high-throughput sequencing (HTS). We obtain 89000242 and 95126160 high-quality clean reads from cDNA libraries of *L.vannamei* with stunted growth symptom (sick group) and health shrimp (health group control), respectively. Totally, 108221 unigenes with an average length of 716 bp were obtained from RNA-seq data. The unigenes were annotated against NR, NT, KO, KOG, SwissProt, PFAM and GO databases. 3975 (3.67%) showed significant matches in all the above databases and 39812 (36.78%) unigenes were annotated in at least one database. Comparative transcriptomic analysis revealed that 807 significantly differentially expressed unigenes were identified, including 282 down-regulated genes and 525 up-regulated genes. Thirteen up-regulated virus-related genes were only discovered in the sick shrimp groups, but not in health ones. Five of them were closely related to virus family Picornaviridae. From phylogenetic tree, we could find five viral genes were closely related to genus Ampvirus, Falcovirus, Aquamavirus (Seal picornavirus). Some up-regulated genes in the Sick groups mainly included genes involved in virus infecting processes concerning antigen processing and presenting, apoptosis, lysosome, phagosome and inflammation. While, many down-regulated genes in the Sick groups mainly involved in the process of starch and sucrose metabolism, and protein digestion and absorption. Our research provided some useful information about pathogenic factor and mechanism on the stunted growth symptom of *L.vannamei*, and was significant for the control and prevention of this disease.

## Introduction

*Litopenaeus vannamei* are economically important and widely farmed aquaculture species. It currently provides about 52% of the total penaeid shrimp production in the world [1–2]. However, the existence of viral or bacterial disease has posed a serious threat to the shrimp aquaculture industry and caused enormous economic losses [3–4]. The stunted growth symptom of *L.vannamei* was one of shrimp diseases appeared frequently in many places in China during the latest several years [5]. Although this significantly impact the shrimp aquaculture and gained considerable attention, the detail for this was not worked out until now.

Previous research indicated that the stunting of Penaeus shrimp was closely related to the presence of some known viruses. Statistical analysis showed a correlation between stunting of *Penaeus monodon* and severity of hepatopancreatic parvovirus (HPV) infection [6]. Rai et al. (2009) detected four kinds of viruses in 72 *P. monodon* samples with slow growth symptom by PCR and suggested the presence of infectious hypodermal and hematopoietic necrosis virus (IHHNV) could be one of the causes of slow growth in cultured *P. monodon* in India [7]. Monodon baculovirus (MBV) was also found a correlation with slow growth of shrimp since the mean length of MBV infected shrimp was significantly shorter than uninfected shrimp from the same pond [8]. In 2001-2002, an unusual retarded growth of *P.monodon*, called monodon slow growth syndrome (MSGS), was found in Thailand. This disease resulted in decreased production of shrimp and approximately US$ 300 million economic losses in 2002 in Thailand [9]. Laem-Singh virus (LSNV) was suggested associating with MSGS [10]. A novel integrase-containing element may interact with LSNV to cause MSGS [11]. RNAi technology using dsRNA of LSNV was proven to be effective in reducing LSNV and restoring normal growth in LSNV-infected shrimp [12].

The specific pathogen free (SPF) shrimp ensure that the *L.vannamei* is free of economically important viruses. This is an important program to ensure the success of shrimp aquaculture. However, new emerging viruses make it very urgent to develop novel and effective method to discover them. In our present work, some *L.vannamei* shrimp samples with stunted growth symptom were collected from Qinzhou and Beihai city in China. The HPV, IHHNV, MBV and LSNV could not be detected in all the samples by PCR test. In order to find out what pathogen agents caused the stunting of shrimp and its pathogenesis, high-throughput sequencing (HTS) technology based RNA-seq and comparative transcriptome analysis were exploited to discover whether new or unusual viruses infected sick shrimp and the molecular mechanism of this disease. Our research may provide some useful information about pathogenic factor and mechanism on the stunted growth symptom of *L.vannamei*, and will be significant for the establishment of more testing standards and maintaining SPF shrimp stocks.

## Materials and Methods

### Shrimp sample

Total of 20 *Litopenaeus vannamei* with symptom of stunted growth (grew for 100 days and the lengths of body were only about 2-3 cm) were collected from shrimps breeding farms (Qinzhou and Beihai city in China), transferred to laboratory immediately and maintained in the aerated recirculating seawater system (temperature 25-26°C, salinity of 32-ppt) for six days prior to the experiment. In parallel, the negative control group was juvenile shrimps collected from a specific pathogen-free (SPF) line with PCR confirmation (without infection of HPV, IHHNV, MBV and LSNV). Then the two shrimp groups: two sick groups with stunted growth (Sick1 and Sick2), and two negative health control groups (each group containing 15 shrimps) (Health1 and Health2) were used in the following experiment. Five individuals from each group were frozen immediately in liquid nitrogen, and then they were stored at −80°C until RNA extraction.

### RNA isolation, cDNA library construction and Illumina sequencing

The above frozen shrimp samples (Sick and Health) were delivered to Novogene Company (Beijing, China) by dry ice for total RNA extraction, cDNA library construction and Illumina sequencing. The total RNA was extracted with TRIzol reagent (Invitrogen, Life Technologies, Grand Island, NY, USA). The RNA purity, concentration, degradation, contamination and integrity was assessed by conventional approach [13–14]. 1.5 µg total RNA was used to generate sequencing libraries by NEBNext ^®^ Ultra™ RNA Library Prep Kit for Illumina ^®^ (NEB, USA). Poly-T oligo-attached magnetic beads were used to isolate mRNA from total RNA. The purified mRNA was cleaved into fragments in Reaction Buffer using divalent cations under elevated temperature. First-strand cDNA was synthesized with random hexamer primer and RNase H, and the second-strand cDNA was produced using RNase H and DNA Polymerase I. Then the double-strand cDNAs were subjected to end repair and addition of A and adapter. The cDNA fragments 150-200 bp in length were selected and purified with AMPure XP system (Beckman Coulter, Beverly, USA). PCR amplification were performed for library construction. After necessary purification and qualification, the cDNA libraries were sequenced on an Illumina Hiseq platform and paired-end reads were generated.

### *De novo* transcriptome assembly and gene function annotation

All the generated raw data (raw reads) were filtered to get high quality clean reads by removing adapter and low quality reads. Meanwhile, Q20, Q30, GC-content and sequence duplication level of the clean data were calculated. Transcriptome assembly was accomplished using Trinity [15] with min_kmer_cov set to 2 by default and all other parameters set default. Gene function of the unigenes was annotated based on databases: Nr (NCBI non-redundant protein sequences); Nt (NCBI non-redundant nucleotide sequences); Pfam (Protein family, http://pfam.xfam.org/search) [16]; KOG/COG (http://www.ncbi.nlm.nih.gov/COG/) [17]; Swiss-Prot (http://www.ebi.ac.uk/uniprot/) and GO (http://www.geneonto-logy.org/). The protein function annotation about Unigenes was achieved by protein with the highest sequence similarity in the databases.

### Differential expression analysis

RSEM was used to estimate the gene expression levels in each sample [18]. The DESeq R package (1.10.1) was used to analyse the different expression pattern between group sick and health. DESeq software package determine differential expression in digital gene expression data by statistical methods. In order to control the false discovery rate (FDR), the P values adjusted and P-value <0.05 was assigned as a criteria for differential expression.

### Enrichment analysis of GO and KEGG

Enrichment analysis of differentially expressed genes (DEGs) was conducted by using Gene Ontology (GO) and KEGG. GO seq R packages based Wallenius non-central hyper-geometric distribution [19] was conducted to adjust for gene length bias in DEGs. The database KEGG was used to understand the gene functions and their activity in cell and organism (http://www.genome.jp/kegg/) [20]. Then the KOBAS software was used to test statistical enrichment of DEGs in KEGG pathways [21].

### Gene expression validation

In order to validate the correction of Illumina sequencing, eight differentially expressed genes were chosen for quantitative analysis by real-time PCR (qRT-PCR) (Table 1). These genes included heat shock 70kDa protein (HSP70), caspase 7 (CASP7), dynein heavy chain 1(DHC-1), low-density lipoprotein receptor (LDLR), trypsin (PRSS), trehalose 6-phosphate synthase (otsA), alpha-amylase (AMY) and collagen (COL1A), where 18s rRNA gene was used as the reference gene. The primers of PCR were designed using Primer Premier 5 software and sequences were indicated in Table 1. Five RNA samples in each group (Sick and Health) were used for gene expression validation. 1 mg of total RNA was used as template for the synthesis of first strand cDNA using reverse transcriptase M-MuLV (NEB). The qRT-PCR reaction mixture (20 μL) contain 2 μL of primer (10 μM), 10 μL 2×SYBR Green PCR buffer, 1 μL of cDNA template. The PCR cycling parameters as follows: 50°C for 2 min, 95°C for 10 min, followed by 40 cycles of 95°C for 20 s, 60°C for 1 min and a final 72°C for 1 min. All the reactions were done in three triplicates.

**Table 1.**
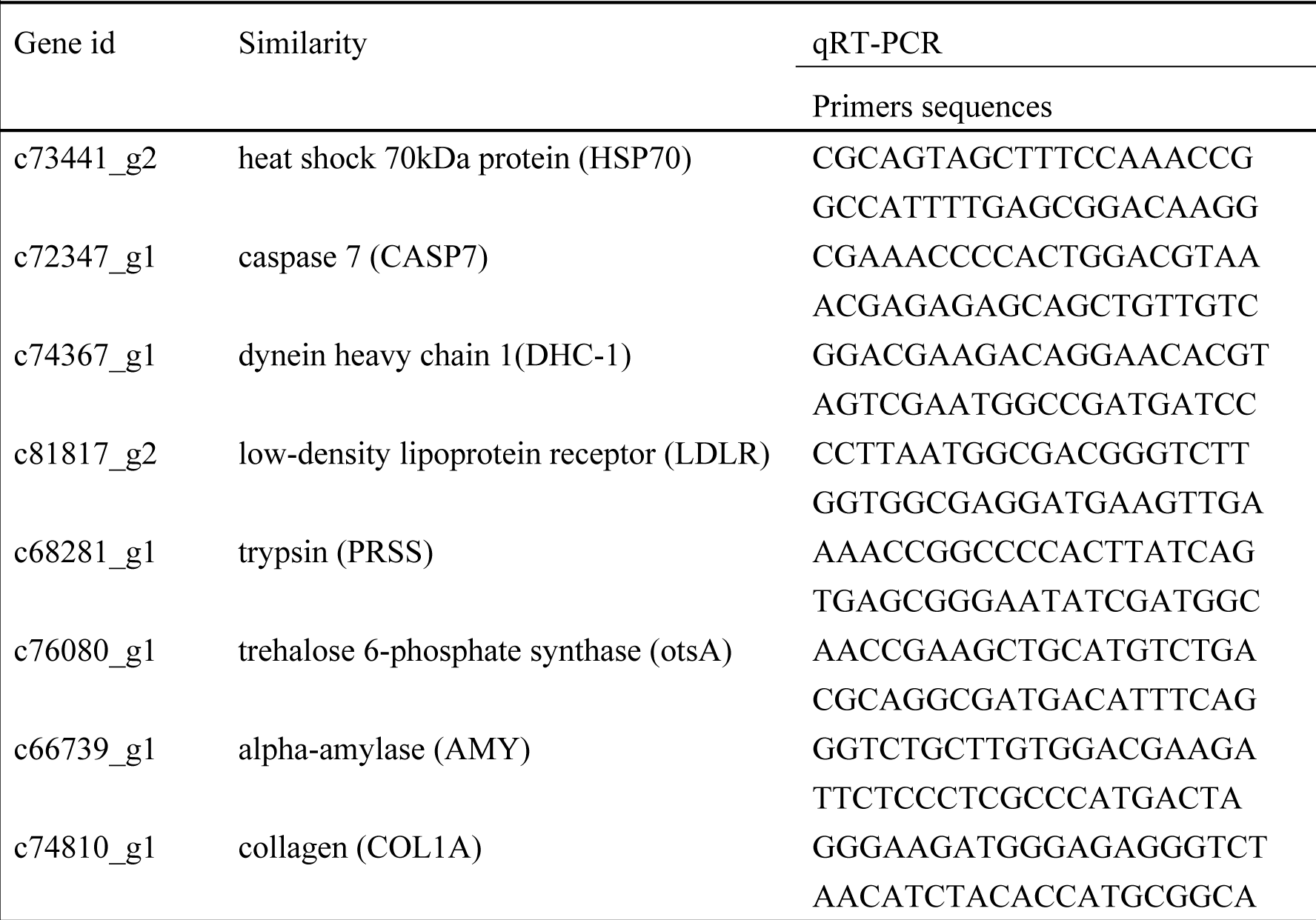
Primers utilized in the gene expression validation experiment.

## Results

### Illumina sequencing and *De novo* assembly of shrimp transcriptome

For the group Sick, 89000242 clean reads were generated from 92277132 raw reads. The mean values of Q20 percentage, Q30 percentage, GC percentage are 96.84%, 92.14%, 49.62%, respectively, and the error rate is 0.02%. For the health samples, 95126160 clean reads were generated from 98615280 raw reads. The Q20 percentage, Q30 percentage, GC percentage are 96.8%, 91.99%, 49.59%, respectively. The clean bases were performed *de novo* assembly using Trinity software. A total of 130544 transcripts with 120846130 nucleotides were assembled. The length of the transcripts ranged from 201 bp to 38462 bp with an average length of 926 bp, and the N50 and N90 were 2080 and 301, respectively. After removing the low-quality and short-length sequences, these transcripts were further assembled and clustered into 108221 unigenes with an average length of 716 bp and N50 of 1361.

### Gene functional annotation

The unigenes were submitted to alignments against the public databases for gene functional annotation, and the unigenes annotated in NR, NT, KO, KOG, SwissProt, PFAM and GO were 26005 (24.02%), 12468 (11.52%), 12194 (11.26%), 16806 (15.52%), 23050 (21.29%), 29084 (26.87%) and 29911 (27.63%), respectively. 3975 (3.67%) showed significant matches in all the above databases and 39812 (36.78%) unigenes were annotated in at least one database.

According to GO analysis, a total of 29912 unigenes were classified into three major functional categories (Fig 1A). The majority kinds of unigenes (1235) involved in the biological process of shrimp cells, followed by 369 kinds of unigenes were responsible for synthesis cellular component. 173 kinds of unigenes had certain molecular function such as catalytic activity, molecular function regulator, transporter activity and binding. In order to determine the biological pathways in both Sick and Health shrimp, all the unigenes were mapped to the referential canonical pathways in the KEGG database. A total of 15205 unigenes were assigned to 229 KEGG pathways. Among them, the top 10 KEGG pathways are: signal transduction (1526), Translation (1438), Transport and catabolism (1194), Endocrine system (1051), protein Folding, sorting and degradation (939), Carbohydrate metabolism (780), Immune system (703), Amino acid metabolism (615), Cellular commiunity (589) and Digestive system (529) (Fig 1B). KOG classification of the unigenes is important for functional annotation and evolutionary studies. A total of 16806 unigenes were categorized into 26 KOG classifications, among which the (R) General Function Prediction Only (2892) was dominant, followed by (O) Posttranslational modification (2389) and (T) Signal transduction mechanisms (2111) (Fig 1C).

**Figure 1.**
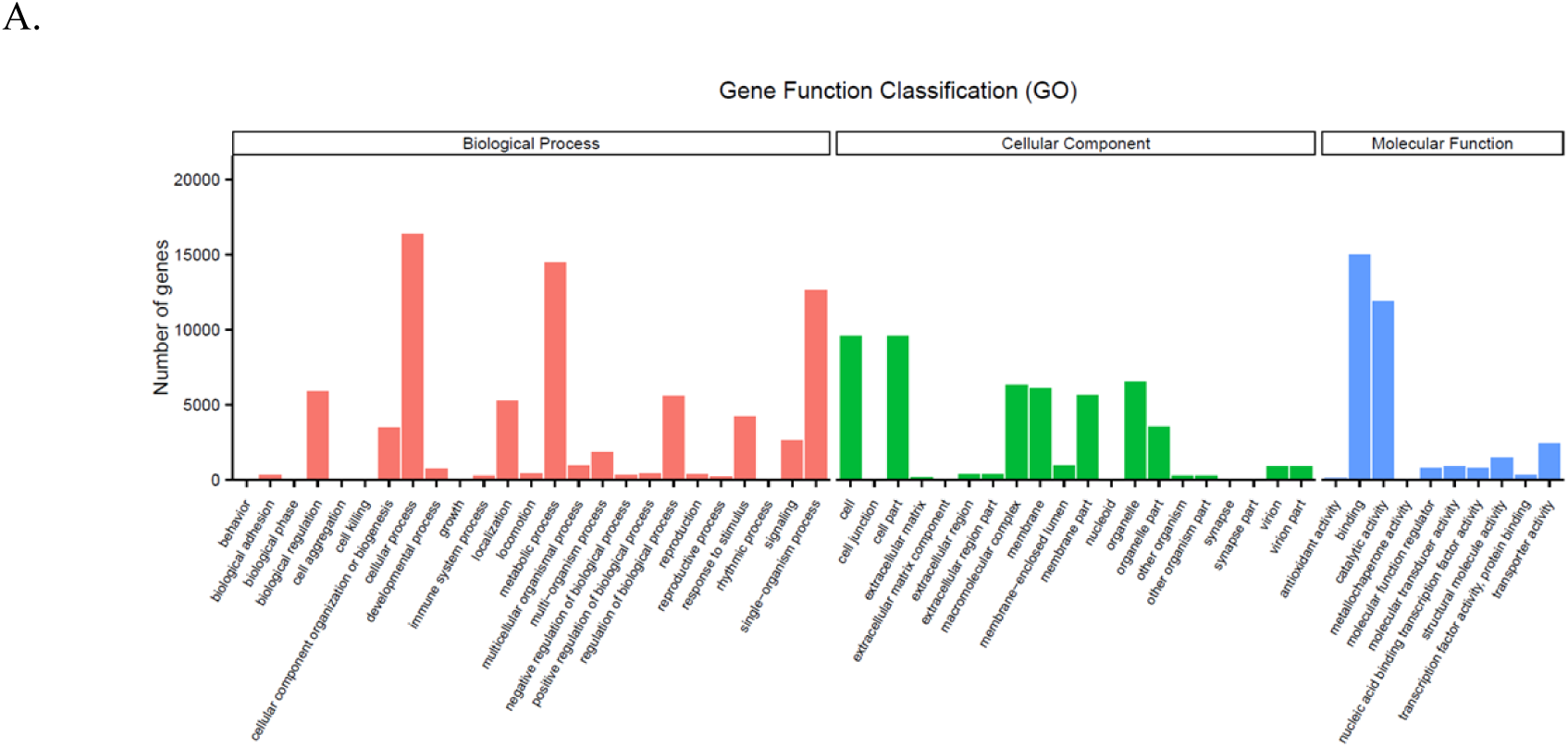
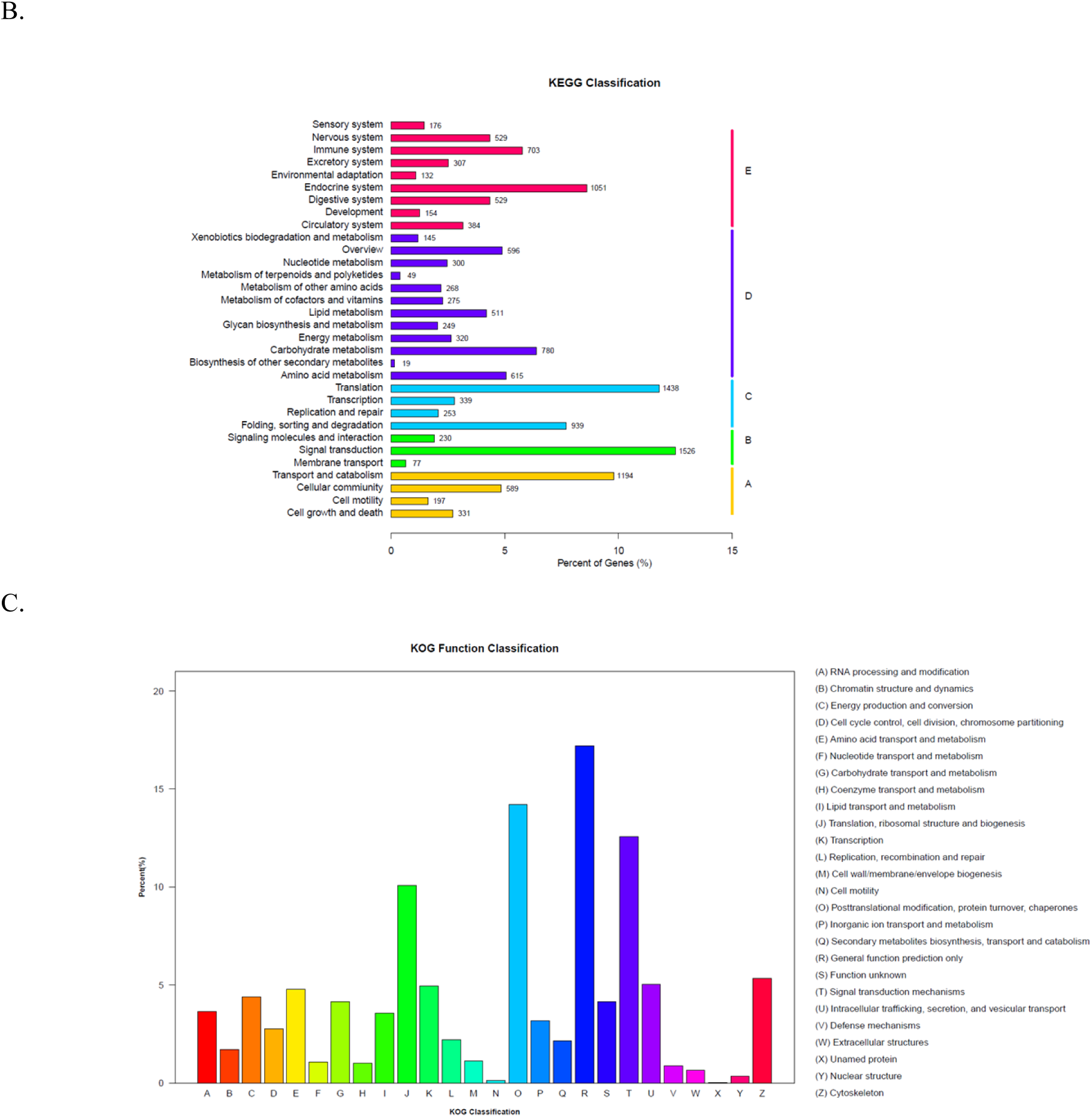
Gene functional annotation by Go, KEGG and KOG analysis. (A) GO categorization of non-redundant unigenes, and each annotated unigene was assigned at least one GO term. (B) KEGG classification of non-redundant unigenes. (C) KOG annotation of putative proteins.

### The Differential expressed genes between Sick and Health shrimp

In our work, an absolute value of log2Ratio≧1 was utilized as the filtering threshold to determine down-regulated or up-regulated genes between Sick and Health shrimp groups. 807 significantly differentially expressed unigenes were identified between Sick and Health groups of shrimp, including 282 down-regulated genes and 525 up-regulated genes (Figure 2). The different expression pattern of Sick and Health were further discovered by a hierarchical cluster analysis. By using this analytical method, a clustered pattern of the DEGs between the two groups was distinctively illustrated (Figure 3), and it was found that the genes expressed in Sick group were distinguishable from that of the Health group.

**Figure 2.**
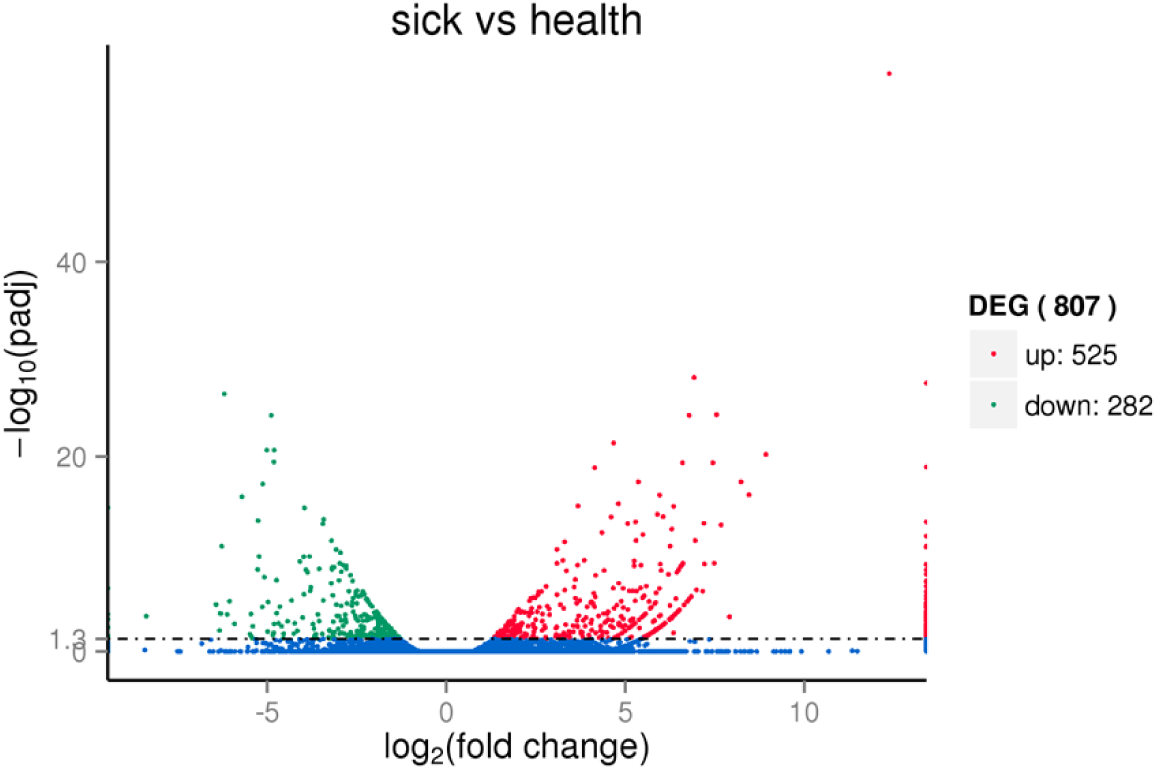
Differentially expressed unigenes were identified between Sick and Health groups of shrimp.

**Figure 3.**
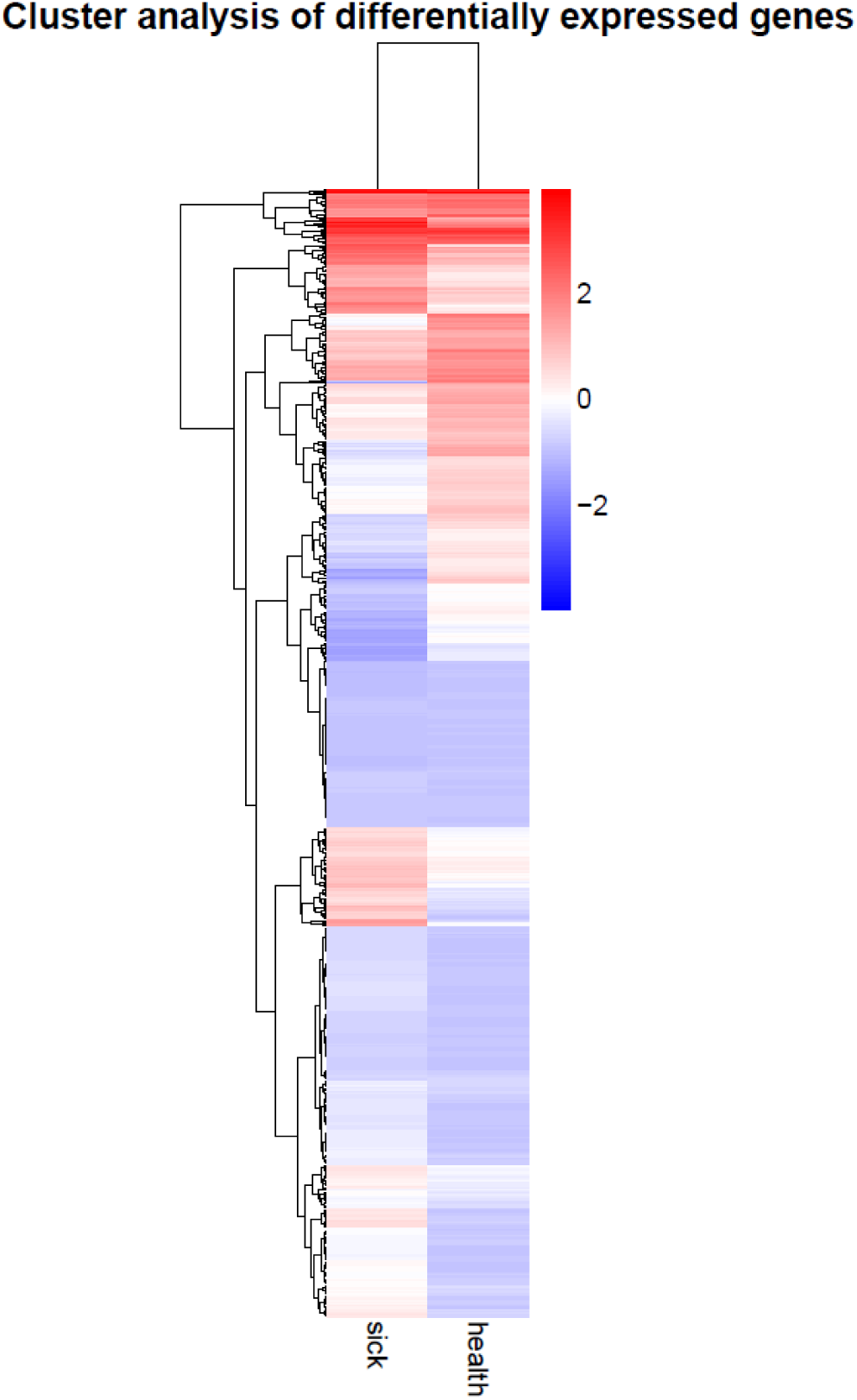
The different expression pattern between Sick and Health groups discovered by a hierarchical cluster analysis.

It was noticed that 35 of 807 DEGs were virus related genes. Among them, 27 viral related genes were up-regulated, while seven were down-regulated (Table 2). It was worth specially mentioning that 13 up-regulated virus-related genes were only discovered in the Sick shrimp groups but not in health ones. Moreover, all the down-regulated virus-related genes appeared in the health shrimp also could be detected in the sick groups. In the 13 up-regulated virus-related genes that only appeared in the sick shrimp groups, it was found that 5 of them were related to genes of virus family Picornavirus (incertae sedis (phylum)/incertae sedis (class)/Picornavirales (order)/Picornaviridae (family)). These viral gene fragments encoded proteins of Picornavirus including RNA-directed RNA polymerase, virus capsid protein and RNA helicase (Table 3). By constructing Neighbour-Joining tree of these five viral gene fragments, it was found that three of them exhibited the same pattern as figure 4A, and the other two displayed trees as figure 4B and 4C, respectively (Figure 4). From all the above phylogenetic tree, we could find a similar feature that the five viral genes were closely related to genus Ampvirus, Falcovirus, Aquamavirus (Seal picornavirus) in the family Picornaviridae. However, the similarity of sequence between the five viruses and that existing in the database was low (about 20%-40%).

**Table 2.**
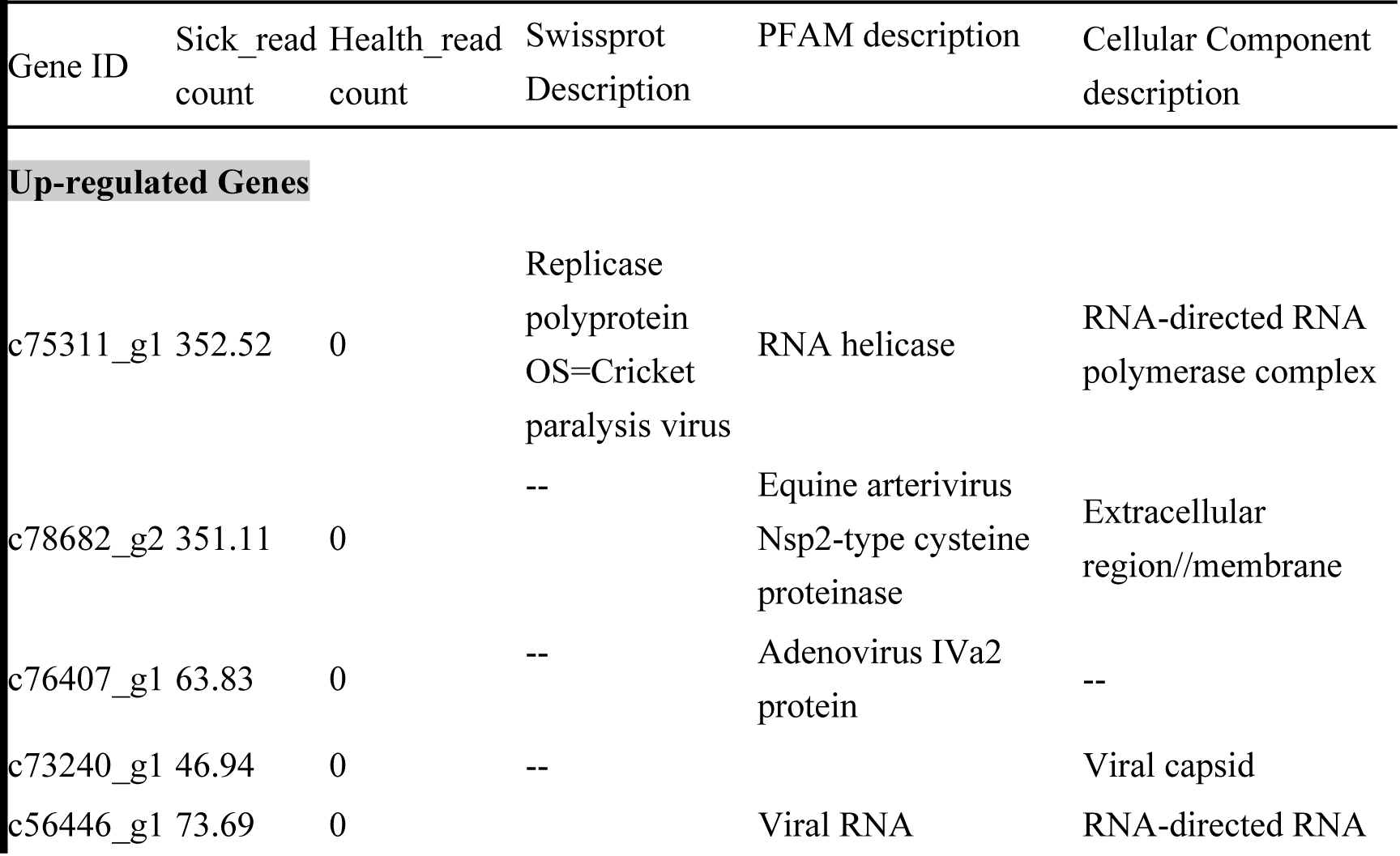
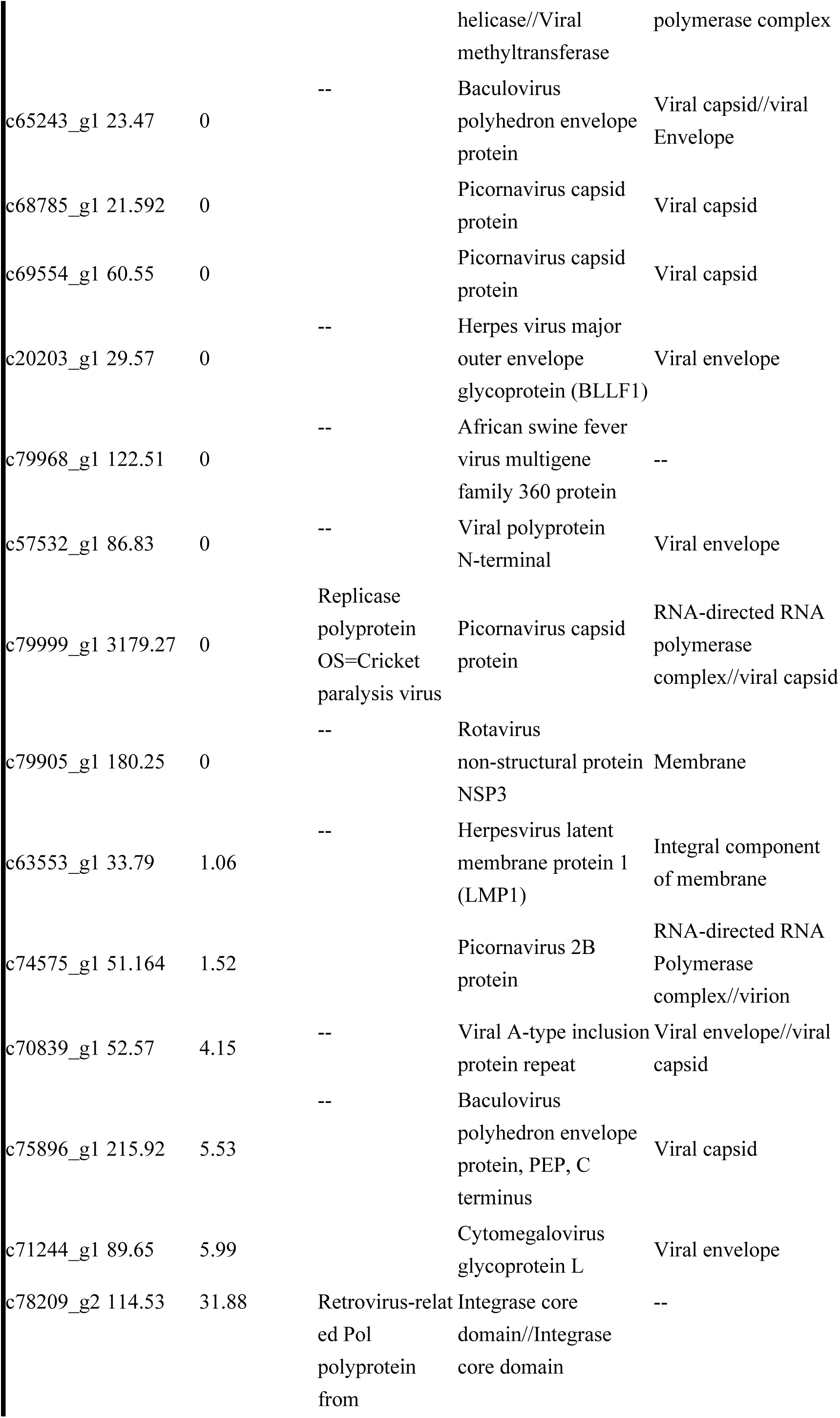
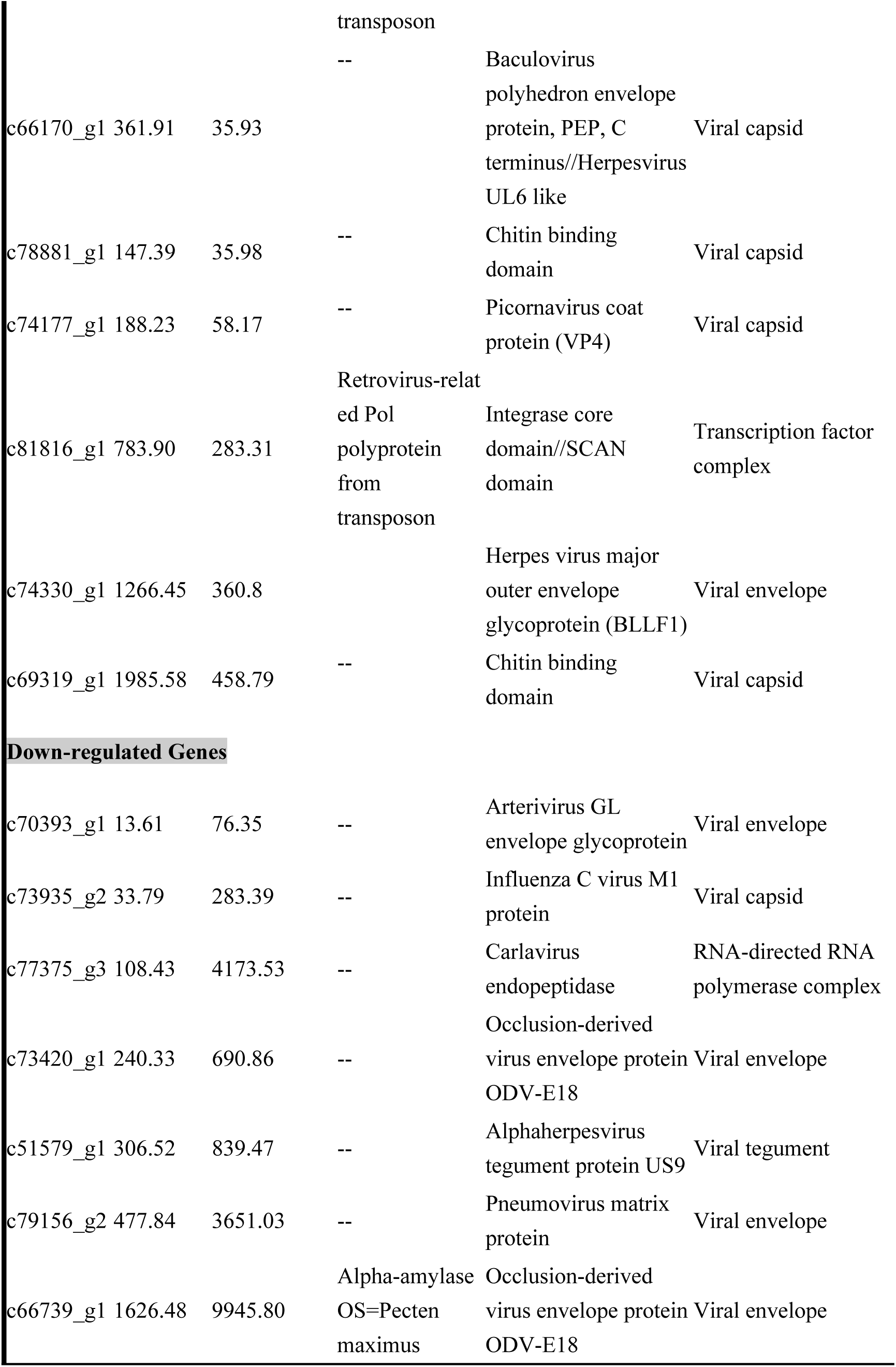
The annotation details about down and up regulated virus-related genes.

**Table 3.** PFAM analysis of five virus-related genes fragments appeared only in the *L.vannamei* groups with stunted growth symptom.

| Fragment number | Length of gene | PFAM prediction |
| --- | --- | --- |
| F1 | 6868 | RNA helicase/RNA-directed RNA polymerase |
| F2 | 4707 | Viral Methytransferase/RNA helicase/RNA-directed RNA polymerase |
| F3 | 1518 | RNA helicase/RNA-directed RNA polymerase/Picornavirus Capsid protein |
| F4 | 4392 | Picornavirus Capsid protein/Dicistro VP4/CRPV Capsid |
| F5 | 9609 | RNA helicase/RNA-directed RNA polymerase |

**Figure 4.**
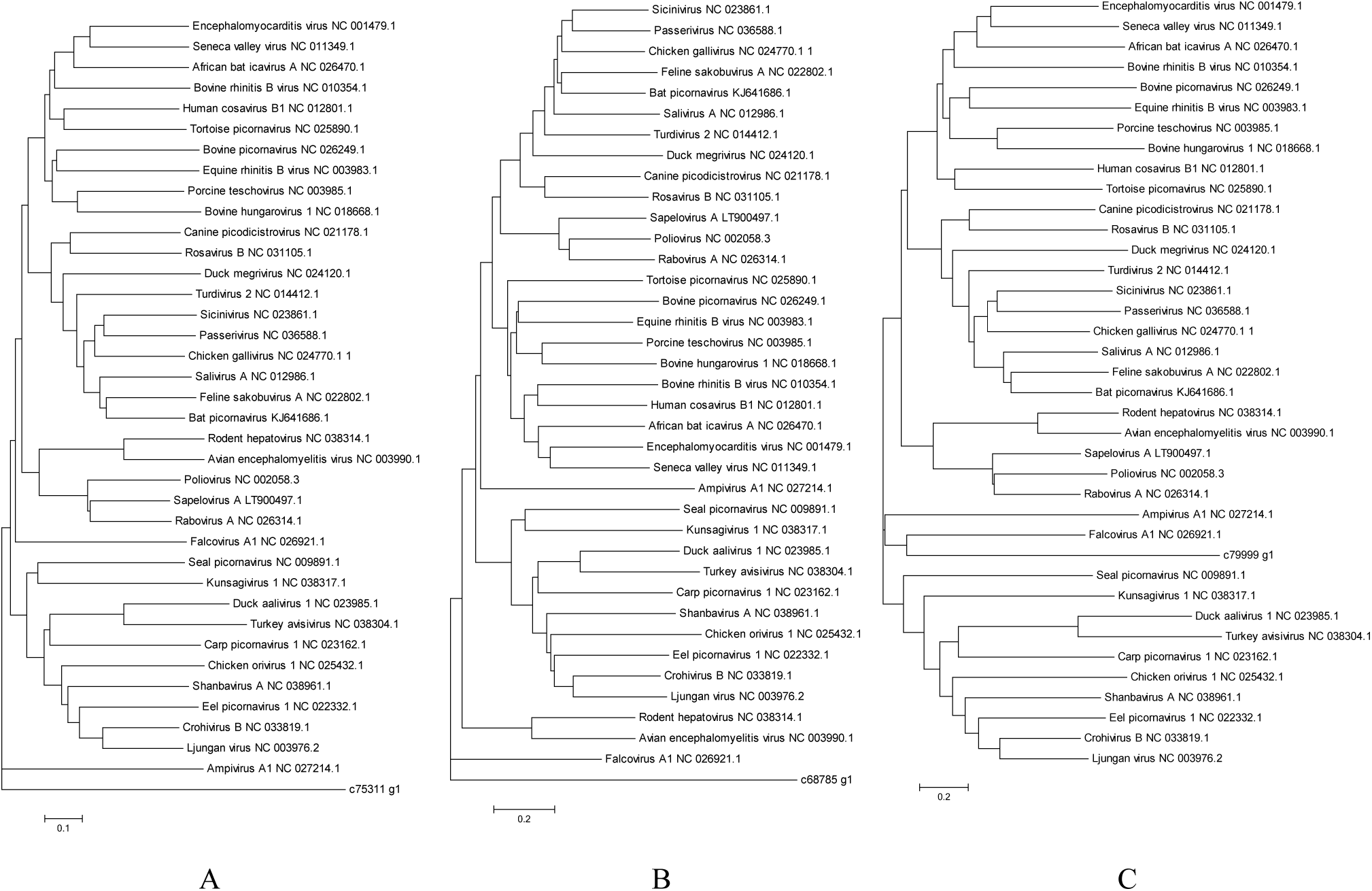
Phylogenetic tree constructed by Neighbour-Joining method using five viral gene fragment sequences and genome sequence of viruses in the family Picornaviridae.

**Figure 5.**
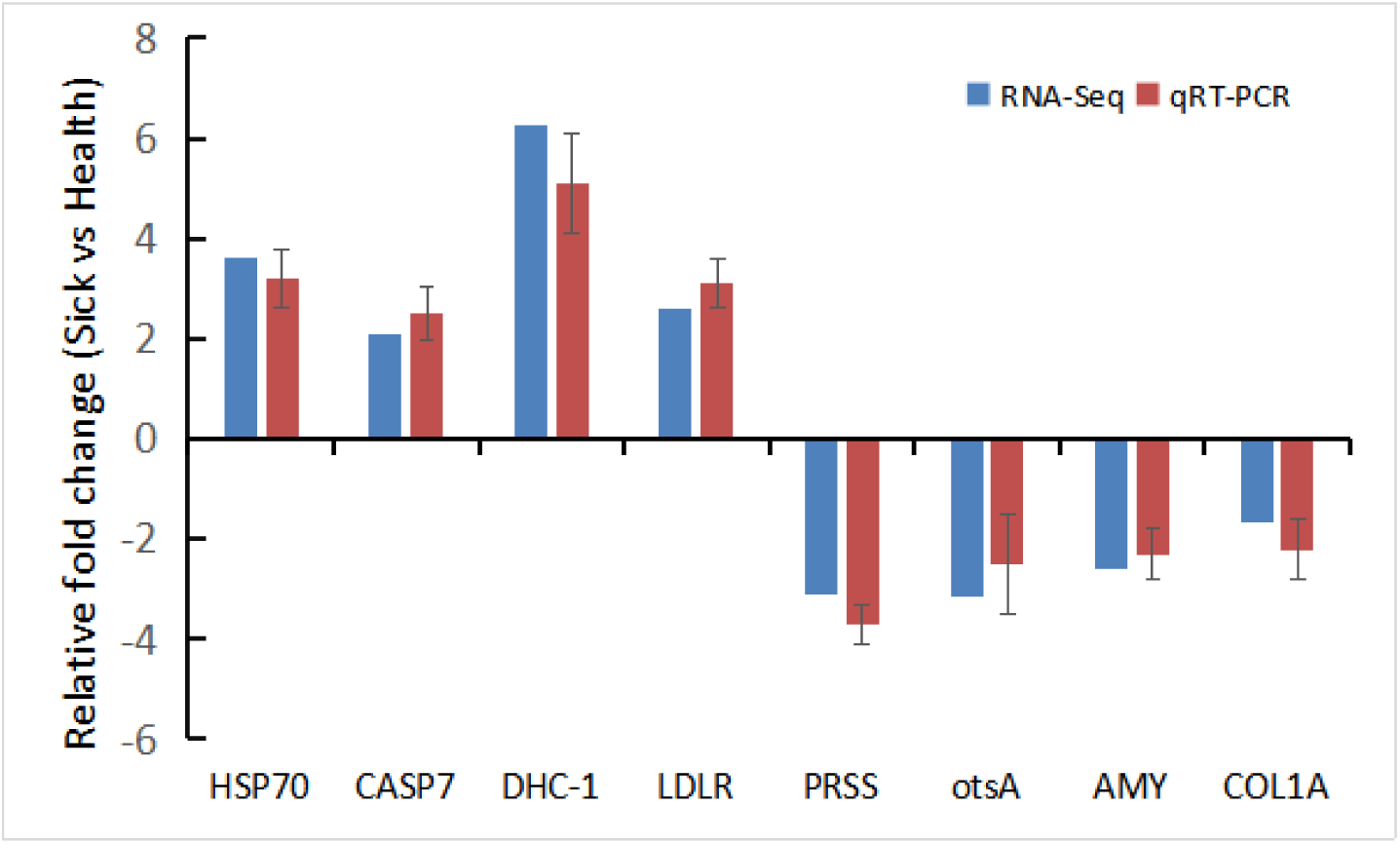
Validation results of RNA-seq profiles by real time PCR. Target gene abbreviations are as follows: HSP 70 - heat shock 70kDa protein, GASP 7 - caspase 7, DHC-1 - dynein heavy chain 1, LDLR - low-density lipoprotein receptor, PRSS - trypsin, otsA - trehalose 6-phosphate synthase, AMY - alpha-amylase, COL1A - collagen.

The DEGs between Sick and Health groups were enriched by KEGG pathway, and a total of 10 significantly changed KEGG pathways (q values < 0.05) were detected (Table 4). Some main up-regulated genes in the Sick groups were concerning biological processes of antigen processing and presenting, apoptosis, lysosome, phagosome, rheumatoid arthritis, systemic lupus erythematosus, and legionellosis. 20 genes enriched in antigen processing and presenting (ko04612) were significantly up regulated, and they encoded HSP 70, HSP 90, legumain and cathepsin B, respectively. 19 genes enriched in lysosome (ko04142) were significantly up regulated, and they encoded clathrin light chain A, lysosomal alpha-mannosidase, legumain and carboxypeptidase C, respectively. 17 genes enriched in phagosome (ko04145) were significantly up regulated, and they encoded actin beta/gamma 1, dynein heavy chain 1, tubulin alpha, tubulin beta and cathepsin L, respectively. 9 genes enriched in rheumatoid arthritis (ko05323) were significantly up regulated, and they all encoded cathepsin L. While, many down-regulated genes in the Sick groups mainly involved in the process of starch and sucrose metabolism, Influenza A infection and protein digestion and absorption. 7 genes enriched in protein digestion and absorption metabolism pathway (ko04974) were significantly down regulated, and they encoded trypsin, neprilysin and collage, respectively. 6 genes enriched in starch and sucrose metabolism (ko00500) were significantly down regulated, and they encoded trehalose 6-phosphate synthase, glucose-6-phosphate isomerase, trehalose 6-phosphate phosphatase, 1,4-alpha-glucan branching enzyme and alpha-amylase, respectively. 8 genes enriched in Influenza A infecting process (ko05164) were significantly down regulated, and they encoded trypsin, HSP 70 and influenza virus NS1A-binding protein, respectively.

**Table 4.**
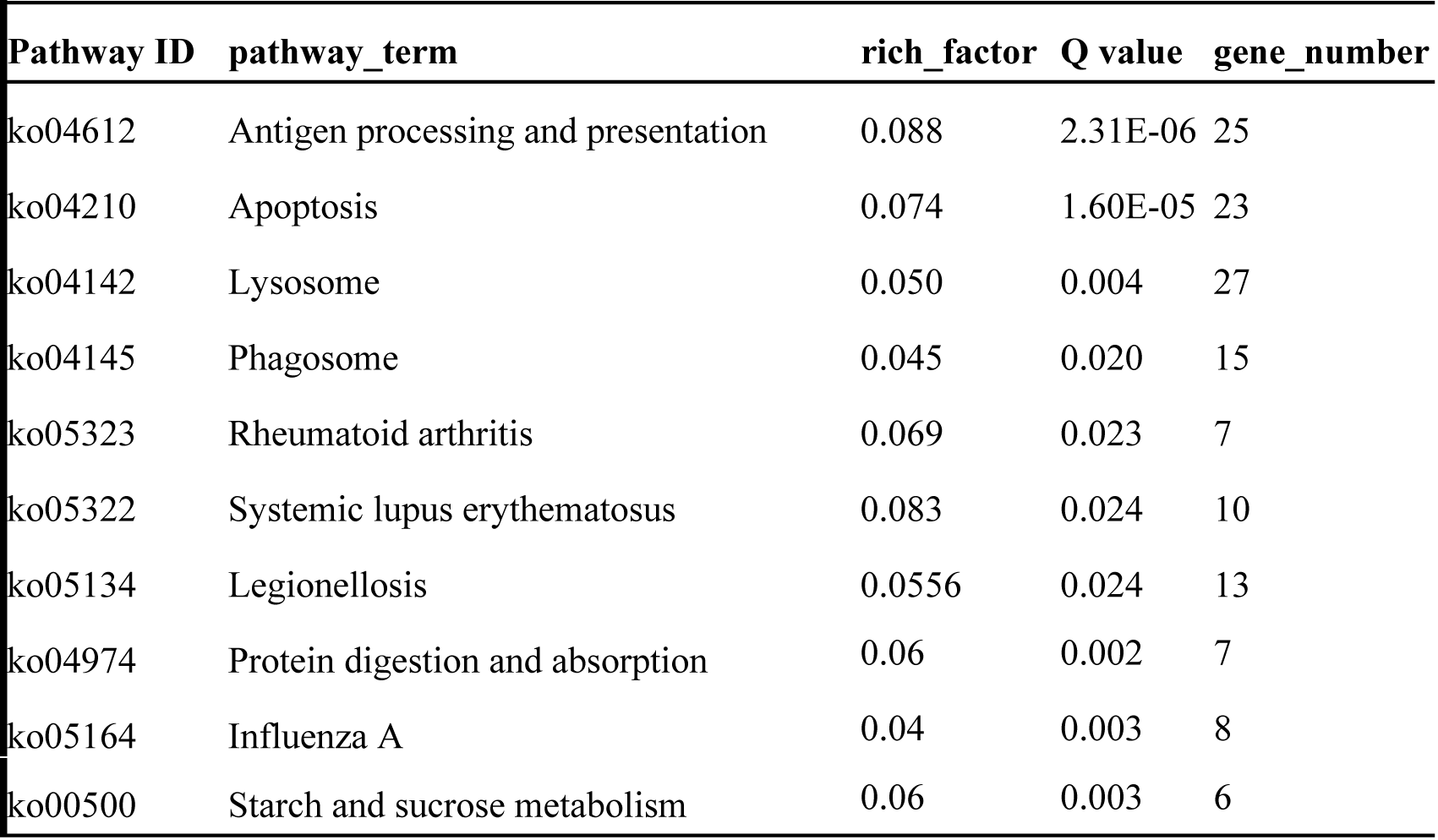
List of 10 significantly changed KEGG pathways (q values < 0.05) in *L.vannamei* with stunted growth symptom.

### Verification of transcriptomic data by qRT-PCR

Eight DEGs were selected for validating the transcriptomic data by real-time PCR analysis. It was found that the quatitive test of up/down-regulated genes indicated by the qRT-PCR and the Illumina sequences demonstrated the similar trends. The qRT-PCR analysis confirmed that transcriptomic data of different expression pattern between Sick and Health shrimp indicated by Illumina sequences were reliable.

## Discussion

Illumina based HTS developed rapidly during the last few years and has been applied abroad in the fields of biological and medical research. It has been employed and proven to be a useful method to discover viruses that are not detected by existing protocols. HTS has been utilized to identify novel plant viruses [22–23]; for instance, HTS was used to find novel virus of the etiology of stem-pitting disease of nectarine [24], viruses associated with decline symptoms of Syrah grapevines [25], apassion fruit woodiness virus isolate from Australia [26]. Such mining of RNA sequence data has also been utilized to identify viruses associated with animal diseases [27–28]. In this study, we demonstrated the potential of the HTS technology platform based comparative transcriptome analysis to identify pathogenic viruses and its pathogenic mechanism involved in stunted growth symptom of *L.vannamei.* We divided the shrimp into two groups: shrimp with stunted growth symptom (Sick group) and shrimp from the normal health pond as negative control (Health group). The total RNA was extracted from intestinal contents and tissue of whole shrimp. By utilized Illumina sequencing, 13 virus gene fragments were only detected in Sick group, but not in Health group. Moreover, the DEGs enriched in stunting shrimp were genes involved in antigen processing and presenting, apoptosis, lysosome, phagosome and inflammation, and many previous research indicated that some of these genes were associated with the process of viral infection. Chen et al. (2013) studied the impact of WSSV infection on host gene expression in the hepatopancreas of *L.vannamei* through the use of RNA-Seq, they discovered some genes involved in the processes of animal defense against pathogens such as apoptosis, antigen processing and presentation and MAPK signaling pathway [29]. It should be mentioned that some genes such as cathepsin B, heat shock protein 70, cathepsin L and legumain were also significantly up regulated in the *L.vannamei* with stunted growth symptom. Li et al. (2013) employed next-generation sequencing (NGS) and bioinformatic techniques to observe the transcriptome differences of the shrimp *Fenneropenaeus chinensis* between latent infection stage and acute infection stage of WSSV [30]. They found many genes involved in acute infection of WSSV, including protein digestion and absorption, phagosome, lysosome, and starch and sucrose metabolism. Coincidentally, these genes were also found in our present work and participated in pathological effect. Similar results were illustrated in other previous research as well [31–34]. Therefore, our obtained data will shed a light on the reason for the occurrence the stunted growth symptom of *L.vannamei* in some place of China, and strongly suggested it may originate from viral infection.

In present work, five Picornavirus (family Picornaviridae) viruses related gene fragments were detected in the stunting *L.vannamei.* Picornaviruses are small non-enveloped viruses, 25-30 nm icosahedral, single messenger-active (+)-RNA, and possess a single long ORF [35]. They could infect and replicate in the cytoplasm of cell, and cause a variety of disease in human and animals [36–37]. Until now, only one member of Picornavirus (family Picornaviridae), Taura syndrome virus (TSV), had been reported in penaeid shrimp [38]. In marine crustacea, only three Picornavirus representatives have been identified, all in crabs [39]. The single strain RNA of TSV genome is 10205 nucleotides in length, containing two large open reading frames (ORFs). The ORF1 represented motifs of a helicase, a protease and an RNA-dependent RNA polymerase; another ORF2 revealed a baculovirus IAP repeat (BIR) domain of inhibitor of apoptosis proteins [40]. In our work, the PFAM analysis indicated that five viruses related genes fragments also contain some conservative domain of Picornavirus genes (e.g. RNA helicase, RNA-directed RNA polymerase and Picornavirus Capsid protein). However, the similarity of sequence between the five viruses and that existing in the database was low (about 20%-40%). Therefore, one interesting result in our work was that some novel Picornavirus may be discovered from *L.vannamei* with stunted growth symptom in China. Their correlation with pathogenesis of stunting of *L.vannamei* require our more in-depth research in the future.

Our another significant finding was that we illustrated some hint about the mechanism of stunting of *L.vannamei* at the molecular level. The fact that viruses infection could result in stunted growth of cell commonly occurred in animal and plant [41–43]. Tang et al. (2016) report that ZIKV virus directly infects human cortical neural progenitor cells with high efficiency, resulting in stunted growth of this cell population and transcriptional dysregulation [44]. Their excellent work solved an urgent global health concern and established a link between ZIKV infection and microcephaly. Qi et al. (2003) discovered a U257A change in the CCR of family Pospiviroidae converted the intermediate strain PSTVdInt to a lethal strain that caused severe growth stunting and premature death of infected plants [45]. Many factor, such as parasite *Enterocytozoon hepatopenaei* (EHP) [46], nutrient elements, and viruses infection could cause slow growth of shrimp. Among them, viruses infection was the most important one and attract widespread attention in the last several years. Although many viral infection (e.g. IHHNV, LSNV, MBV, HPV) were discovered leading to the slow or stunting growth of Penaeus shrimp in previous research, the mechanism has been not illustrated until now [6–8]. In our study, the result that some viruses were only detected in *L.vannamei* with stunted growth symptom made us hypothesize that this disease may have correlation with viral infection. This hypothesis was further supported by gene expression profiles, since many genes of shrimp, which participate in the infecting process of viruses, were up-regulated in sick shrimp with stunting symptom. Moreover, we have found the reason why *L.vannamei* infected by viruses could not grow up to the normal size. This may originated from the reason that the shrimp infected by viruses could not metabolize protein and sugar (starch and sucrose) very well. The correlation between viral infection and metabolism of protein and sugar has been well studied in plant [47–50]. However, there were not any information about this in animal. Anyway, the depressed metabolism of protein and sugar could undoubtedly inhibit the growth of cell in plant or animal [51–53].

Although our research provided some useful information about pathogenic factor and mechanism on the stunted growth symptom of *L.vannamei*, more work should be carried out in the future. HTS based comparative transcriptome analysis was a helpful method for discovering novel viruses infecting shrimp, but what can not be neglected, this method was more effective in detecting RNA viruses than DNA viruses. We are not sure about whether DNA viruses are significant in the occurrence of this shrimp disease. Thus, metagenomic assay will be utilized for further work on the identification and proven of other pathogenic factor [54–55]. Moreover, the cDNA probe and RT-PCR methods will be useful tools for detecting Picornavirus in more sick shrimp samples and its correlation with the emerging of stunted growth symptom. Anyway, our work was important for the establishment of more testing standards and maintain SPF shrimp stocks.

## Acknowlege

Supported by the National Natural Science Foundation of China (31560727); Guangxi Natural Science Foundation (2014GXNSFBA118135, 2018JJA130187).

